# Mutations in *HOMEOBOX DOMAIN-2* improve grain protein content in wheat without significantly affecting grain yield and senescence

**DOI:** 10.1101/2025.10.10.681749

**Authors:** Luqman B. Safdar, Ian D. Fisk, Marianna Pasquariello, Aneesh Lale, Iain R. Searle, Rahul A. Bhosale, M. John Foulkes, Scott A. Boden

**Affiliations:** School of Agriculture, Food and Wine, University of Adelaide, Glen Osmond, SA 5064, Australia; School of Biosciences, University of Nottingham, Loughborough, LE12 5RD, United Kingdom; International Flavour Research Centre, Division of Food Nutrition and Dietetics, University of Nottingham, Loughborough, LE12 5RD, United Kingdom; International Flavour Research Centre (Adelaide), School of Agriculture, Food and Wine and Waite Research Institute, University of Adelaide, Glen Osmond, SA 5064, Australia; Department of Crop Genetics, John Innes Centre, Norwich Research Park, Norwich, NR4 7UH, United Kingdom; Council for Agricultural Research and Economics, Research Centre for Genomics and Bioinformatics, Via San Protaso 69, Fiorenzuola d’ Arda (PC), 29017, Italy; School of Biological Science, University of Adelaide, Adelaide, SA 5005, Australia

**Keywords:** wheat, yield, senescence, vasculature, grain protein, breeding, *HOMEOBOX DOMAIN-2*

## Abstract

MicroRNA-resistant alleles of *HOMEOBOX DOMAIN-2* (*HB-2*) were recently reported to improve grain protein content (GPC) in wheat by modifying vascular development in stems to increase amino acid distribution to grain. However, it is not known if these alleles increase GPC at the expense of other physiologically important traits, such as senescence and yield-component traits. Here, we evaluated the effects of mutations in *HB-2* on grain yield and nitrogen (N)-use efficiency as well as senescence and plant architecture in multi-environment glasshouse and field experiments. In glasshouse experiments, the mutations in *HB-2* moderately improved GPC irrespective of N supply and without accelerating senescence or decreasing grain number or weight compared to the wildtype spring wheat cv. Cadenza. The trait analysis indicated mutant plants were more tolerant of N limitation in terms of above-ground N reduction compared to the wild type. The expression of all *HB-2* homeologues increased in spikes at the lemma primordium stage in response to N limitation, indicating the later GPC increase could be a result of enhanced N sink strength determined during early development, rather than processes caused by direct changes in *HB-2* expression in the grain. In field experiments, the introgression of *HB-2* alleles into Australian elite spring wheat cultivars significantly improved GPC without compromising grain yield. The outcomes of this study support the introduction of miRNA-resistant *HB-2* alleles, with a preference to *HB-A2*, into commercial wheat for improving grain quality without a yield penalty.

## INTRODUCTION

Recently, mutations in a homeobox-domain leucine zipper class III transcription factor *HOMEOBOX DOMAIN-2* (*HB-2*) were reported to increase grain protein content (GPC) in bread wheat by altering vascular architecture **(Dixon *et al*., 2022)**. The identified mutations in *HB-2* on the A and D sub-genomes (termed *HB-A2* and *HB-D2*, respectively) locate to the complementary sites for microRNA165/166 (miR165/166), which disrupt cleavage of *HB-2* transcripts and increase its expression. The higher *HB-2* expression promotes formation of additional vascular bundles in the stem, which facilitates greater translocation of amino acids, or N, to sink organs, such as the spike and grain. It was proposed that the increased N supply helps boost GPC. Importantly, the higher GPC was reported to occur without affecting grain weight **(Dixon *et al*., 2022)**. This highlights the miR165/166 resistant *HB-2* alleles as strong candidates for incorporation into breeding programs aimed at augmenting grain quality. A major constraint in improving GPC is its frequent association with a yield penalty, and research on many fronts is attempting to disrupt this trade-off: for example, by nutrient and soil management, selecting for grain protein deviation (GPD), breeding for *Gpc-B1* functional alleles, and identifying new alleles from unexplored germplasm **(Safdar *et al*., 2023b)**. The *Gpc-B1* functional allele improves GPC by accelerating senescence, which remobilises more N from source tissue to grain after anthesis **(Uauy *et al*., 2006b)**. The accelerated senescence reduces chlorophyll content in leaves and disrupts photosynthesis, which can lead to yield losses **(Tabbita *et al*., 2017)**. Senescence has been a widely targeted mechanism for genetic improvement of GPC for nearly two decades.

Higher protein improves bread-making quality of wheat as it is associated with enhanced gluten levels, which improve dough rheology. GPC is also an economically important trait in some countries where grain that contains protein above a threshold of ca. 13% is sold at a premium **(Barak *et al*., 2013)**. The discovery of modified plant vasculature induced by mutations in *HB-2* offers an alternate approach for the genetic improvement of GPC, potentially without yield compromises. Modifying plant vasculature to achieve optimum grain quality also has the potential to reduce N_2_O emission from agricultural soil by reducing the use of N fertiliser **(Safdar *et al*., 2023a)**. It is, however, critical to fully evaluate the impact of the *HB-2* alleles on yield components, N accumulation and N-remobilisation efficiency, which were not evaluated previously. *HB-2* alleles were also associated with secondary spikelet development, which could affect overall thousand grain weight (TGW) and/or the fruiting efficiency (i.e., grains per g spike dry matter at anthesis) **(Dixon *et al*., 2022)**. Similarly, it is crucial to measure other key physiological traits, such as leaf senescence, a developmental process that involves expressional changes in thousands of genes that ultimately impact grain protein, yield and N-use efficiency **(Sultana *et al*., 2021)**. Senescence is strongly linked to plant N dynamics as N is a key component of chlorophyll structure and Rubisco, and is therefore crucial for photosynthesis, biomass and yield **(Fageria *et al*., 2010)**. Plants that can utilise N efficiently, by maintaining grain quality characteristics under reduced N inputs, will be desirable for wheat breeding.

This study was designed to examine the effects of miR165/166 resistant *HB-2* alleles on physiological traits of plants grown under different N conditions. This was done by analysing traits related to inflorescence architecture, N-use efficiency components, yield components, GPC, leaf senescence and photosynthesis. Ultimately, the goal of discovering beneficial alleles is to introduce them into elite commercial cultivars so that improved traits can be passed to breeders, growers and consumers. The enhanced expression of *HB-2* improved GPC due to increased uptake and assimilation of amino acids from the stem into the inflorescence because the vascular architecture was modified **(Dixon *et al*., 2022)**. However, N uptake per plant was not reported previously. This study aimed to test the hypothesis that the higher *HB-2* expression increases N availability in the stem and spike during early developmental stages, allowing the grains of mutant genotypes to accumulate more protein.

## MATERIALS AND METHODS

### Glasshouse experiment - University of Nottingham

The first experiment evaluated the phenotypic performance of wheat TILLING lines carrying miR165/166 resistant *HB-2* alleles and their wild type siblings in optimal N and N-stress environments. Germplasm derived from two lines (*CAD1290* and *CAD1761*), which carry mutations in *HB-D2* and *HB-A2*, respectively, were screened for GPC prior to the start of the experiments (see Plant material and sowing). GPC levels were confirmed using hyperspectral imaging (HSI) to confirm genotypic differences before sowing. HSI-based protein determination was derived from an in-house single grain protein calibration. The experiment was conducted at the Sutton Bonington Campus, 52°49’57.4“N 1°15’00.1“W, from January to June 2022. Sowing date was 17 January, and natural light was extended to 16 h photoperiod till 21 March and then kept to natural daylength. The temperature was maintained at 20 °C before transplanting and then the plants were kept frost free. The experimental design used was a split-plot where two levels of N treatment (HN: high N and LN: low N) were randomised on the main-treatment and genotypes were randomised on the sub-plot treatment.

### Plant material and sowing

Germplasm used for this experiment was from a backcrossed generation (BC_3_F_4_) derived from two lines *CAD1290* and *CAD1761* of the Cadenza TILLING population (JIC, UK) **(Krasileva *et al*., 2017; Dixon *et al*., 2022)**. *CAD1290* contains a mutation in the miR165/166 binding site of *HB-D2* and *CAD1761* has a near-identical mutation in *HB-A2. CAD1290* lines included three genotypes: 1290 wild type (1290WT), heterozygous (1290HT) and mutant (1290MT). *CAD1761* lines included 2 genotypes: 1761 wild type (1761WT) and mutant (1761MT), with the “wild-type” being a homozygous control with the contains the reference allele derived from the back-crosses **(Dixon *et al*., 2022)**. The 1290 MT was initially included, but was subsequently removed because it displayed reduced plant stature and curled leaf phenotypes that prevent accurate comparison of grain yield traits **(Dixon *et al*., 2022)**. The dominant allele of *HB-D2* induces strong upregulation in 1290HT, and so this genotype was considered the variant genotype. KASP markers were used to accurately segregate WT, HT and MT genotypes from 1290HT **(Dixon *et al*., 2022)**.

DNA was extracted from leaf tissues using the cetyltrimethylammonium bromide (CTAB) method **(Yu *et al*., 2017)**. To confirm *CAD1290* genotypes, KASP analysis was performed as described previously **(Dixon *et al*., 2022)**. Genotypes were confirmed using two allele specific KASP assays **(Supplementary Table 1)** and further validated using Sanger sequencing of HB-D2 specific amplicons **(Supplementary Fig. 1; Supplementary Table 2)**.

### Transplanting into the glasshouse and experimental design

40 plants per genotype were transplanted into 2 L pots (one plant per pot), which included five plants (technical replicates) per replicate, with four biological replicates sown per genotype. The pots were filled with low N compost (Klasmann medium peat) and irrigated with full nutrient feed minus N; 357 pots including 200 for experimental plants and 150 for border plants **(Supplementary Fig. 2)**. Automatic irrigation was set up with complete feed minus N. N was applied separately from NH_4_NO_3_ (34% N) in two doses for low N and three doses for high N. Low N dose corresponded to 60 kg N/ha and high N dose to 200 kg N/ha that is the optimum N level for wheat fields in the UK. Low N dose was split into two doses of 30 kg N/ha and high N into three doses of 100, 50 and 50 kg N/ha. The first dose was applied at transplantation, the second at growth stage (GS) 31 and third at GS39 only for high N **(Zadoks *et al*., 1974)**. Two fungicide applications were applied to all pots to control powdery mildew – a sulphur evaporator was also installed in the glasshouse. The N calculation for the amount of NH_4_NO_3_ to add per pot for the N treatments was based on 1 g NH_4_NO_3_ = 0.34 g N and the pot diameter of 15 cm. A TinyTag data logger was installed in the glasshouse and the daily minimum and maximum temperature were recorded (no extreme temperatures above 32 °C were observed) **(Supplementary Fig. 3)**.

### Phenotyping for GPC, spike architecture and yield components

GPC was measured at 15 and 30 days after anthesis from developing grain and at harvest using the Dumas Combustion method, GPC analysis of grain grown at University of Adelaide from the TILLING lines elite backcrossed lines was performed using the Elementar Rapid N Exceed Nitrogen and Protein Analyser (Elementar). The plant architecture and yield related traits that were measured included: primary and secondary spikelets per spike (SpS) and fertile and infertile SpS at anthesis and harvest, spike length (SL) and rachis length (RchL), grain weight (GW) and number (GN) distribution in the top, middle and bottom sections of the main-shoot spike (dividing by fertile spikelet number and rounding up). We also measured final leaf number (LN) of the main shoot and plant height (PH).

The number of fertile (FT, those with a spike) and infertile tillers (IFT) at anthesis and harvest was counted; and TGW, GN, chaff dry weight (ChW), rachis weight (RcW), grain weight (GY) and harvest index (HI) were determined for the main shoot and fertile tillers separately at harvest. Dry matter (DM) was recorded after drying for 48 h at 80 °C; and dry matter partitioning was determined at anthesis and harvest that included lamina, stem (including leaf sheath) and spike (grain and chaff at harvest) components from the main shoot and tillers.

### Determination of senescence kinetics

Flag-leaf chlorophyll concentration was determined from heading till maturity using a handheld SPAD meter **(Konica Minolta SPAD-502Plus)**. SPAD readings were fitted against days after heading (DAH, GS55) using the Gompertz curve method in **python-3.9.7** to determine the timing of the inflection point (IP) and the rate of change at IP. The Gompertz equations was:

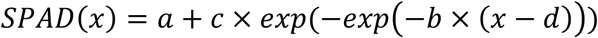

Where: *a* is asymptote representing the final SPAD value after senescence, *b* is the rate of change at the inflection point, *c* is the difference between the initial SPAD value and asymptote and *d* is inflection point, when the maximum senescence rate occurs. For each replicate of each G × N combination, parameters were calculated using ***curve_fit*** from ***scipy*** library for nonlinear least-squares fitting **(Virtanen *et al*., 2020)**. Parameters *d* and *b* representing the timing of the inflection point and the rate of change at the inflection point were extracted for statistical analysis.

To determine the photosynthetic performance of plants, flag-leaf gas exchange data were recorded using a LI-6400/XT Portable Photosynthesis System (Licor Biosciences, Lincoln, NE, USA). The Li-Cor 6400 settings were: flow rate 400 µmol s^−1^, block temperature 20°C with ambient relative humidity. The sample (cuvette) CO_2_ concentration was set to 400 µmol mol^− 1^ and PAR was set to 1500 µmol m^−2^ s^−1^ (10% blue). The leaf temperature and the leaf transpiration rate were measured. The Fv’/Fm’, which is the maximum proportion of light energy which PSII is capable of using for its photochemistry: its “maximum quantum efficiency”, was also measured via the chlorophyll fluorescence readings from the Li-Cor 6400. One plant per genotype was harvested at anthesis for dry matter partitioning and N partitioning. The stem, spike and leaf samples from the main shoot were transported to Campden BRI (Chipping Campden, UK) for determination of total N using the Kjeldahl method. Three plants per genotype were used for GPC analysis of developing grains. The 5th plant (on which the flag-leaf SPAD and gas-exchange measurements were taken) was grown till maturity and harvested for analysing grain yield components and N-related traits as described above. N% was determined from main shoot leaf and stem (including sheath) at maturity using Kjeldahl.

### N partitioning and N-remobilisation efficiency

To understand the effect of the mutations on N allocation to grain sink and on N-use efficiency (NUE), we analysed a broad range of N-related traits including: above-ground N (AGN) and N partitioning index (NPI) for plant components at anthesis and harvest, post-anthesis N uptake (PANU), post-anthesis N remobilisation (NR) from stem and leaf to the grain, NR efficiency (NRE), N-uptake efficiency (NUpE), N-utilisation efficiency (NUtE) and NUE **(Gaju *et al*., 2014; Gaju *et al*., 2011)**. AGN was determined by combining the N uptake from different plant components. NPI was determined as the ratio of the N in the plant component to the total AGN, for example Leaf-NPI = Leaf N / AGN and so on for Stem- and Spike-NPI at anthesis and Leaf-, Stem-, Chaff-, Rachis- and Grain-NPI at harvest. PANU was determined as the amount of AGN at harvest that was absent in the above-ground plant at anthesis. NR was determined as the amount of N in the plant component at anthesis that was absent in the plant component at harvest. NRE was determined based on the calculation **(Gaju *et al*., 2014)**:

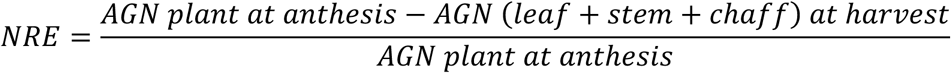

NUpE was determined as the ratio of N uptake to N supply, NUtE as the ratio of GY to N uptake by main shoot and NUE as the ratio of GY to total N supply. Nitrogen harvest index (NHI) was determined as the ratio of grain N to AGN harvest. All analyses were performed on the main shoot.

### Statistical analysis

A split-plot analysis of variance (ANOVA) model was fitted in **R-4.4.1** using ***agricolae*** package **(de Mendiburu, 2019)**. Post-hoc analysis was conducted using Fisher’s LSD method with Benjamini-Hochberg adjusted *p* value for controlling the false discovery rate. Means and standard deviations were calculated using ***dplyr*** package **(Wickham *et al*., 2021)**. The results were plotted using ***ggplot2* (Wickham, 2016)**. Two analyses were designed for *CAD1290* and *CAD1761*, with each including two N levels (High N and Low N) as whole plot factors, two genotypes (1290WT and 1290HT or 1761WT and 1761MT) as subplot factors, and four replicates (Blocks) as random factors. The model equation was:

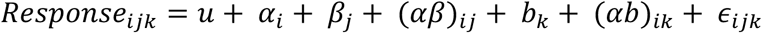

Where: *u* is overall mean, *⍺*_*i*_ is fixed effect of *i*th genotype, *β_j_* is fixed effect of *j*th treatment, (*⍺β*)_*ij*_ is interaction between genotype and treatment, *b*_*k*_ is random effect of *k*th block, (*⍺b*)_*ik*_ is random effect of genotype within a block (whole plot error) and *∈*_*ijk*_ is residual (subplot error).

Genotype and N level were considered as main effects and their interaction G × N was also assessed; significance was set at α = 0.05. Comparisons between N levels within each genotype were conducted using the ***emmeans*** package with estimated marginal means compared for significant difference **(Lenth *et al*., 2021)**.

### Glasshouse experiment at University of Adelaide

Two further glasshouse experiments were conducted to analyse senescence and *HB-2* expression under different N levels at The Plant Accelerator (TPA, University of Adelaide, Australia. For each experiment, daylength and maximum daily temperature were artificially maintained to 16 h and 22 °C, respectively. In the first experiment, *CAD1290* and *CAD1761* lines were grown under three N levels of 80, 160 and 240 kg N/ha (equivalent) to investigate senescence timing and GPC. *CAD1761* seedlings were grown immediately in pots whereas *CAD1290* seedlings were genotyped to select WT and HT genotypes using KASP, as described earlier, before being transplanted to 2L pots with no N cocopeat soil medium. The experiment was designed in a random complete block design with five replicates per genotype **(Supplementary Fig. 4)**. The source of N application for this experiment was urea, and the calculations were made by calculating the soil area per pot with reference to the N levels (equivalents) per ha.

### *HB-2* expression analysis via RT-qPCR

The second experiment focused on N-responsiveness of *HB-2* expression in wild-type Cadenza. Grains were planted under normal growth conditions in a full nutrient soil minus N. The N doses (based on urea) were applied separately in two phases. For each N level (80, 160 and 240 kg N/ha), the total N dose was split into two. The first dose (1/2 total N) was applied at transplantation (into 2L pots, no N cocopeat soil medium) one-week post-germination, and the second (2/2 N) applied two weeks later. Immature spike and subtending stem tissue were collected at the lemma primordium (LP) and terminal spikelet (TS) stages. Developing spikes from five plants/replicate, immature stems from three plants/replicate, and young and mature leaf tissues from one plant/replicate were collected, with sampling between 1000-1400.

RNA was extracted using the Spectrum Total RNA Extraction Kit as per the manufacturer’s instructions (Sigma-Aldrich). RNA was reverse transcribed into cDNA using the SuperScript^TM^ III kit (Thermo Fisher Scientific) as per the manufacturer’s instructions. RT-qPCR was performed on four biological replicates of each sample in a 7900HT Fast Real-Time PCR system (Applied Biosystems) using GoTaq qPCR Master Mix (Promega). All RT-qPCR data points were the average of two technical replicates, and three biological replicates. Expression of candidate genes was normalised against *ACTIN* and *GAPDH*.

### Analysing the effect of *HB-2* alleles on GPC and yield in elite wheat backgrounds

To determine the effect of microRNA-resistant *HB-2* alleles in elite wheat cultivars, we introduced the mutant *HB-A2* (*CAD1761*) and *HB-D2* (*CAD1290*) alleles into three Australian elite cultivars, Mace, Rockstar and Sherrif through marker-assisted selection until the BC_2_F_4_ generation. These crosses produced the following genotypic groups: Mace/1761WT and Mace/1761MT, Rockstar/1761WT and Rockstar/1761MT, and Sherrif/1761WT and Sherrif/1761MT; Mace/1290WT, Mace/1290HT and Mace/1290MT, Rockstar/1290WT, Rockstar/1290HT and Rockstar/1290MT, and Sherrif/1290WT, Sherrif/1290HT and Sherrif/1290MT. Trials were conducted at two experimental field sites: University of Adelaide, Roseworthy Campus 34°31’37.9“S 138°41’21.0“E and Australian Grain Technologies, 34°32’07.4“S 138°45’41.8“E. The sites contained loam over red clay soils, and the trials were sown in early May 2023, with nitrogen applied at 100 kg/ha across two intervals (seeding and early tillering). Data for *CAD1761* crosses yielding two genotypes (WT and MT) were analysed by independent samples *t-*test. Data for *CAD1290* crosses yielding three genotypes were analysed by one-way ANOVA, followed by Tukey’s HSD for testing significant differences in means. G × E interaction was analysed by a two-way ANOVA. All analyses were done using ***SciPy*** and ***statsmodels*** libraries in **python-3.9.7 (Seabold and Perktold, 2010; Virtanen *et al*., 2020)**.

## RESULTS

### Genotyping and hyperspectral imaging confirm TILLING lines with higher GPC

The experiments measuring grain protein, yield and associated traits at Nottingham and Adelaide will hereafter be referred to as Experiment 1 (Expt. 1) and Experiment 2 (Expt. 2), respectively. Before starting Expt. 1, protein levels were determined non-destructively in grains of TILLING lines that formed modified spikelet architecture, including lines with miR165/166 resistant *HB-2* alleles, to select genotypes for further analysis. Hyperspectral images of grains were examined to collect spectral profiles, and GPC was predicted using an in-house supervised learning model (International Flavour Research Centre, University of Nottingham). Consistent with previous results, the predicted GPC values in mutants were higher than the wild type lines **(Supplementary Fig. 5)**. Overall, the predicted GPC values were relatively high, in the range 13-16%, possibly because of the training data used in predictive model not having information from the TILLING population; nonetheless, the observed trends were as expected. Based on these preliminary results, *CAD1290* and *CAD1761* were selected to investigate the effect of the mutations in *HB-2* on plants. For *CAD1290*, the heterozygous genotype was selected as the variant, rather than the homozygous mutant, because it displays strong upregulation of *HB-D2* without the detrimental leaf curling and short stature traits of the homozygous mutant (see Materials and Methods).

### *HB-2* alleles moderately enhance GPC without a significant yield loss

To test if the GPC enhancement by miR165/166 resistant *HB-2* alleles depends on soil N availability, GPC was analysed in developing grain at 15 days after anthesis (DAA) and at harvest in Expt. 1. The lines carrying the resistant *HB-2* alleles (1290HT and 1761MT) increased GPC regardless of the N level (i.e., 200 and 60 kg N/ha–eq.), although the increase at harvest was non-significant **(Table 1)**. In 1290HT, GPC was significantly higher compared to WT at 15 DAA under optimum N supply and moderately increased under low N. GPC difference at 15 DAA for 1290HT compared to 1290WT was +16.81% under high N and +2.37% under low N; and for 1761MT compared to 1761WT was −2.51% under high N and +4.62% under low N **(Table 1)**. GPC difference at harvest for 1290HT compared to 1290WT was +0.52% under high N and +4.23% under low N; for 1761MT compared to 1761WT it was +10.42% under high N and −0.32% under low N **(Table 1)**. Although the *HB-2* alleles in both mutant genotypes showed GPC improvement, the timing of GPC accumulation was unique to each line; 1290HT accumulated more GPC during early development, which became diluted towards maturity, whereas 1761MT apparently accumulated more GPC near maturity.

**Table 1:** Split-plot analysis of variance for GPC and yield components in experiment 1. Values in table represent mean ± standard deviation (n = 4). Percent change refers to change in phenotypes from wild type to mutant.

| Genotype | High N | Low N | Difference | ANOVA |
| --- | --- | --- | --- | --- |
| **Grain protein content (%) 15 DAA** |  |  |  |  |
| 1290W | 11.3±0.3 | 10.13±0.32 | 1.17 | N: p = 0.0012** |
| 1290H | 13.2±0.46 | 10.37±0.25 | 2.83 | G: p = 0.0146* |
| Pct. Change | 16.81% | 2.37% | | G×N: p = 0.0259* |
| 1761W | 11.97±0.87 | 10.83±0.4 | 1.13 | N: p = 0.2085 |
| 1761M | 11.67±0.91 | 11.33±0.72 | 0.33 | G: p = 0.7922 |
| Pct. Change | -2.51% | 4.62% | | G×N: p = 0.4598 |
| **Grain protein content (%) at harvest** |  |  |  |  |
| 1290W | 9.65±0.26 | 8.52±0.24 | 1.13 | N: p = 0.0164* |
| 1290H | 9.7±0.68 | 8.88±1.02 | 0.82 | G: p = 0.6194 |
| Pct. Change | 0.52% | 4.23% | | G×N: p = 0.63 |
| 1761W | 9.6±0.79 | 9.35±0.44 | 0.25 | N: p = 0.3098 |
| 1761M | 10.6±2.58 | 9.32±0.15 | 1.28 | G: p = 0.4834 |
| Pct. Change | 10.42% | -0.32% | | G×N: p = 0.4841 |
| **Thousand grain weight (g) 15 DAA** |  |  |  |  |
| 1290W | 19.58±2.89 | 16.46±0.36 | 3.12 | N: p = 0.1429 |
| 1290H | 18.96±2.6 | 18.12±0 | 0.83 | G: p = 0.7805 |
| Pct. Change | -3.17% | 10.09% | | G×N: p = 0.3515 |
| 1761W | 17.5±2.86 | 16.25±2.86 | 1.25 | N: p = 0.0314* |
| 1761M | 20.42±1.57 | 15.21±0.36 | 5.21 | G: p = 0.5743 |
| Pct. Change | 16.69% | -6.40% | | G×N: p = 0.1172 |
| **Thousand grain weight (g) at harvest** |  |  |  |  |
| 1290W | 59.73±8.37 | 55.43±3.97 | 4.3 | N: p = 0.1252 |
| 1290H | 48.06±4.46 | 41.9±5.09 | 6.16 | G: p = 0.0405* |
| Pct. Change | -19.54% | -24.41% | | G×N: p = 0.7623 |
| 1761W | 51.12±0.86 | 49.69±7.15 | 1.43 | N: p = 0.6946 |
| 1761M | 45.24±14.68 | 43.26±3.03 | 1.97 | G: p = 0.2767 |
| Pct. Change | -11.50% | -12.94% | | G×N: p = 0.9499 |
| **Grain number from main shoot** |  |  |  |  |
| 1290W | 64±18.02 | 59±13.24 | 5 | N: p = 0.748 |
| 1290H | 68.5±20.34 | 68±14.24 | 0.5 | G: p = 0.5816 |
| Pct. Change | 7.03% | 15.25% | | G×N: p = 0.7923 |
| 1761W | 59±5.03 | 50.25±9 | 8.75 | N: p = 0.2273 |
| 1761M | 69±4.08 | 69.25±7.8 | -0.25 | G: p = 0.0582. |
| Pct. Change | 16.95% | 37.81% |  |  |
| **Grain yield (g) from main shoot** |  |  |  |  |
| 1290W | 3.71±0.62 | 3.23±0.51 | 0.48 | N: p = 0.214 |
| 1290H | 3.29±0.98 | 2.82±0.51 | 0.47 | G: p = 0.328 |
| Pct. Change | -11.32% | -12.69% | | G×N: p = 0.9888 |
| 1761W | 3.02±0.28 | 2.46±0.33 | 0.56 | N: p = 0.1589 |
| 1761M | 3.13±1.05 | 3±0.39 | 0.13 | G: p = 0.5472 |
| Pct. Change | 3.64% | 21.95% | | G×N: p = 0.3601 |
| **Main shoot dry matter (g)** |  |  |  |  |
| 1290W | 8.16±1.14 | 7.57±1.15 | 0.59 | N: p = 0.4294 |
| 1290H | 7.15±1.77 | 6.62±0.97 | 0.54 | G: p = 0.2787 |
| Pct. Change | -12.37% | -12.55% | | G×N: p = 0.9673 |
| 1761W | 7.19±0.34 | 6.2±0.29 | 0.99 | N: p = 0.0042** |
| 1761M | 7.27±1.21 | 6.86±0.74 | 0.41 | G: p = 0.6021 |
| Pct. Change | 1.11% | 10.65% |  | G×N: p = 0.1151 |
| **Main shoot fertile spikelet per spike at harvest** |  |  |  |  |
| 1290W | 18.75±3.2 | 20±4.16 | -1.25 | N: p = 0.4061 |
| 1290H | 19±3.65 | 20.5±2.08 | -1.5 | G: p = 0.8806 |
| Pct. Change | 1.33% | 2.50% |  | G×N: p = 0.9379 |
| 1761W | 19.25±1.26 | 19.25±1.26 | 0 | N: p = 1 |
| 1761M | 19.25±1.5 | 19.25±1.5 | 0 | G: p = 1 |
| Pct. Change | 0 | 0 |  | G×N: p = 1 |
| **Fertile tillers at harvest** |  |  |  |  |
| 1290W | 4.75±1.26 | 2.25±0.5 | 2.5 | N: p = 0.0027** |
| 1290H | 5.5±1 | 2.5±0.58 | 3 | G: p = 0.2522 |
| Pct. Change | 15.79% | 11.11% |  | G×N: p = 0.6704 |
| 1761W | 6.25±0.5 | 2.5±0.58 | 3.75 | N: p = 0*** |
| 1761M | 5.25±0.5 | 2.25±0.5 | 3 | G: p = 0.0796. |
| Pct. Change | -16% | -10% |  | G×N: p = 0.1682 |
| **Grain yield (g) from tillers** |  |  |  |  |
| 1290W | 13.46±2.31 | 4.01±1.65 | 9.45 | N: p = 2e-04*** |
| 1290H | 10.63±1.46 | 3.66±0.49 | 6.98 | G: p = 0.0595. |
| Pct. Change | -21.03% | -8.73% |  | G×N: p = 0.2614 |
| 1761W | 13.93±1.26 | 4.12±1.3 | 9.8 | N: p = 4e-04*** |
| 1761M | 10.28±3.67 | 3.18±0.9 | 7.1 | G: p = 0.0877. |
| Pct. Change | -26.20% | -22.82% |  | G×N: p = 0.3088 |
| **Tillers dry matter (g)** |  |  |  |  |
| 1290W | 30.72±5.57 | 11.16±2.64 | 19.55 | N: p = 2e-04*** |
| 1290H | 26.74±3.23 | 10.09±1.28 | 16.65 | G: p = 0.1893 |
| Pct. Change | -12.96% | -9.59% |  | G×N: p = 0.5307 |
| 1761W | 33.46±2.21 | 11.38±2.08 | 22.08 | N: p = 0*** |
| 1761M | 27.44±2.67 | 8.72±1.22 | 18.72 | G: p = 0.0162* |
| Pct. Change | -17.99% | -23.37% |  | G×N: p = 0.1922 |
| **Grain yield (g) per plant** |  |  |  |  |
| 1290W | 17.17±2.3 | 7.24±1.6 | 9.93 | N: p = 1e-04*** |
| 1290H | 13.92±2.27 | 6.47±0.76 | 7.45 | G: p = 0.1052 |
| Pct. Change | -18.93% | -10.64% |  | G×N: p = 0.2578 |
| 1761W | 16.94±1.37 | 6.58±1.35 | 10.36 | N: p = 7e-04*** |
| 1761M | 13.4±4.7 | 6.18±0.87 | 7.23 | G: p = 0.251 |
| Pct. Change | -20.91% | -6.10% |  | G×N: p = 0.298 |
| **Harvest index (%)** |  |  |  |  |
| 1290W | 44.68±2.58 | 38.9±5.22 | 5.79 | N: p = 0.0202* |
| 1290H | 42.61±2.64 | 39.43±3.62 | 3.18 | G: p = 0.6043 |
| Pct. Change | -4.63% | 1.36% |  | G×N: p = 0.3971 |
| 1761W | 41.78±1.82 | 37.65±4.12 | 4.13 | N: p = 0.6069 |
| 1761M | 39.35±9.93 | 39.86±4.43 | -0.51 | G: p = 0.9754 |
| Pct. Change | -5.82% | 5.87% |  | G×N: p = 0.5136 |

Significance codes: '\*\*\*' 0.001 '\*\*' 0.01 '\*' 0.05

The overall observed GPC increase was lower than previously reported, although the trend was similar. The previous data presented by **Dixon *et al*. (2022)** was NIR based; similar results were seen in HSI based GPC prediction in our study (see **Supplementary Fig. 5**), which suggests spectral based GPC predictions tend to be higher compared to the N measurement-based ground-truth analysis. To validate the effect of miR165/166 resistant *HB-2* alleles on GPC, all four genotypes were re-evaluated in Expt. 2; similar results were obtained with moderate GPC increase under different N levels in the genotypes with miRNA-resistant *HB-2* alleles **(Supplementary Fig. 6)**. To test the effect of these *HB-2* alleles on yield component traits, we analysed thousand grain weight at corresponding stages. TGW reduced at harvest in mutant lines under both N levels, with a significant decrease observed for 1290HT compared to 1290WT (*p* 0.0405*; **Table 1**). The reduction in TGW could be complemented by higher grain number in lines with mutant *HB-2* alleles, caused by paired spikelet formation. Grain number (GN) increase was observed for both genotypes with mutant alleles (1290HT and 1761MT), compared to their wild type siblings **(Table 1)**. Higher GN per spike is considered a direct indicator of high yields in wheat under optimum conditions, as it is associated with higher post-anthesis grain sink demand for assimilate **(Reynolds *et al*., 2022)**.

Grain yield (GY) per main shoot was slightly lower in 1290HT compared to 1290WT, although the difference was non-significant **(Table 1)**. There was no difference in the main shoot fertile spikelets per spike (SpS) at harvest in either *CAD1290* or *CAD1761* genotypes. Main-shoot DM showed no effect in either family; however, the trend was downwards in 1290HT (−12.37% under low N and −12.55% under high N) and upwards in 1761MT (+1.11% under low N and +10.65% under hight N) compared to wild types **(Table 1)**. Moreover, there was no difference in the fertile tillers per plant at harvest in the case of *CAD1290* genotypes, and tiller GY was moderately reduced (*p* 0.0595). In *CAD1761*, the mutant had higher (although non-significant) GY per main shoot under both N treatments compared to the wild type. The fertile tillers per plant at harvest were moderately reduced in 1761MT compared to 1761WT (*p* 0.0796), which reduced GY of tillers moderately (*p* 0.0877) and DM yield significantly (*p* 0.0162*) in the mutant **(Table 1)**. The overall decrease in GY was moderate and non-significant, with more yield per tiller in 1761MT. All other yield components showed no effect of genotype or G × N interaction terms.

The dry matter (DM) partitioning at anthesis revealed that under N limitation, main-shoot lamina DM (%) decreased in all genotypes. The main-shoot stem DM (%) increased in all genotypes but the ratio of increase was much smaller in 1761MT (0.58%) compared to other genotypes (∼6%); a significant genotype effect (*p* 0.0323*) and G × N interaction (*p* 0.0349*) was observed for *CAD1761* genotypes **(Supplementary Table 3)**. Contrastingly, main-shoot spike DM (%) was reduced under N limitation in 1761WT but increased in 1761MT (genotype effect for *CAD1761 p* 0.0086*; **Supplementary Table 3**). This showed that under low N, 1761MT partitioned relatively less biomass to the stem compared to WT and relatively more to the spike. Trends for similar effects were observed at harvest **(Supplementary Table 4)**. These findings suggested that relatively more resources were allocated to the spike in the case of 1761MT (*HB-A2* allele) at low N compared to the wild-type. Our results showed that both GY and above-ground DM were unaffected in *CAD1290* by the miR165/166 resistant *HB-2* alleles. Similarly, both GY and DM yield per main shoot were unaffected in 1761MT irrespective of the N level; in fact, both showed an increasing (but non-significant) trend, compared to WT. However, the reduction in tillers per plant in 1761MT may have a negative effect on grain yield (t/ha), which should be assessed in future trials. These results also point to differences in phenotypes following mutation of *HB-D2*, relative to *HB-A2,* consistent with previous reports about leaf curling traits in these lines **(Dixon *et al*., 2022; Jiang *et al*., 2023)**.

### MicroRNA resistant *HB-2* alleles do not accelerate the onset of senescence

Leaf senescence is a well-known mechanism involved in regulating protein accumulation through N-remobilisation to grain **(Sultana *et al*., 2021)**. However, since there is evidence *HB-2* alleles improved N-remobilisation through modifications in plant vasculature in previous work, we hypothesised that *HB-2* does not accelerate the timing of senescence. To test this hypothesis, we investigated flag-leaf chlorophyll content over developmental time after anthesis. This was achieved by fitting flag-leaf SPAD readings against days after heading to determine the time taken by a leaf to reach the onset of senescence (taken as the inflection point; when SPAD is fixed at 36.8% of the upper asymptote) and the rate at which chlorophyll content declined at the inflection point **(Supplementary Fig. 7)**. Analysis of the inflection point timing showed slightly accelerated senescence in 1290HT (non-significant) and a delayed senescence in 1761MT (*p* 0.0424*) compared to their wild types. The genotype effect in *CAD1761* was only observed under high N (G × N *p* 0.0205*). In other words, instead of earlier senescence, 1761MT took seven more days to reach the onset of senescence compared to 1761WT under optimum N supply, whereas no difference was observed under low N **(Fig. 1A; Supplementary Table 5)**. Moreover, under optimum N, the rate of change of leaf chlorophyll content at the inflection point – although not reaching the significance threshold – was considerably slower for 1761MT (−0.11±0.03 d^−1^) compared to 1761WT (− 0.26±0.12 d^−1^; **Fig. 1B; Supplementary Table 5**). This means that the mutant not only reached the onset of senescence later but also senesced more slowly under optimum N. These results indicate that neither the onset of senescence nor rate of decline in chlorophyll content was accelerated by the miR165/166 resistant *HB-2* alleles. The only observed change was in 1761MT, where senescence was delayed.

**Figure 1:**
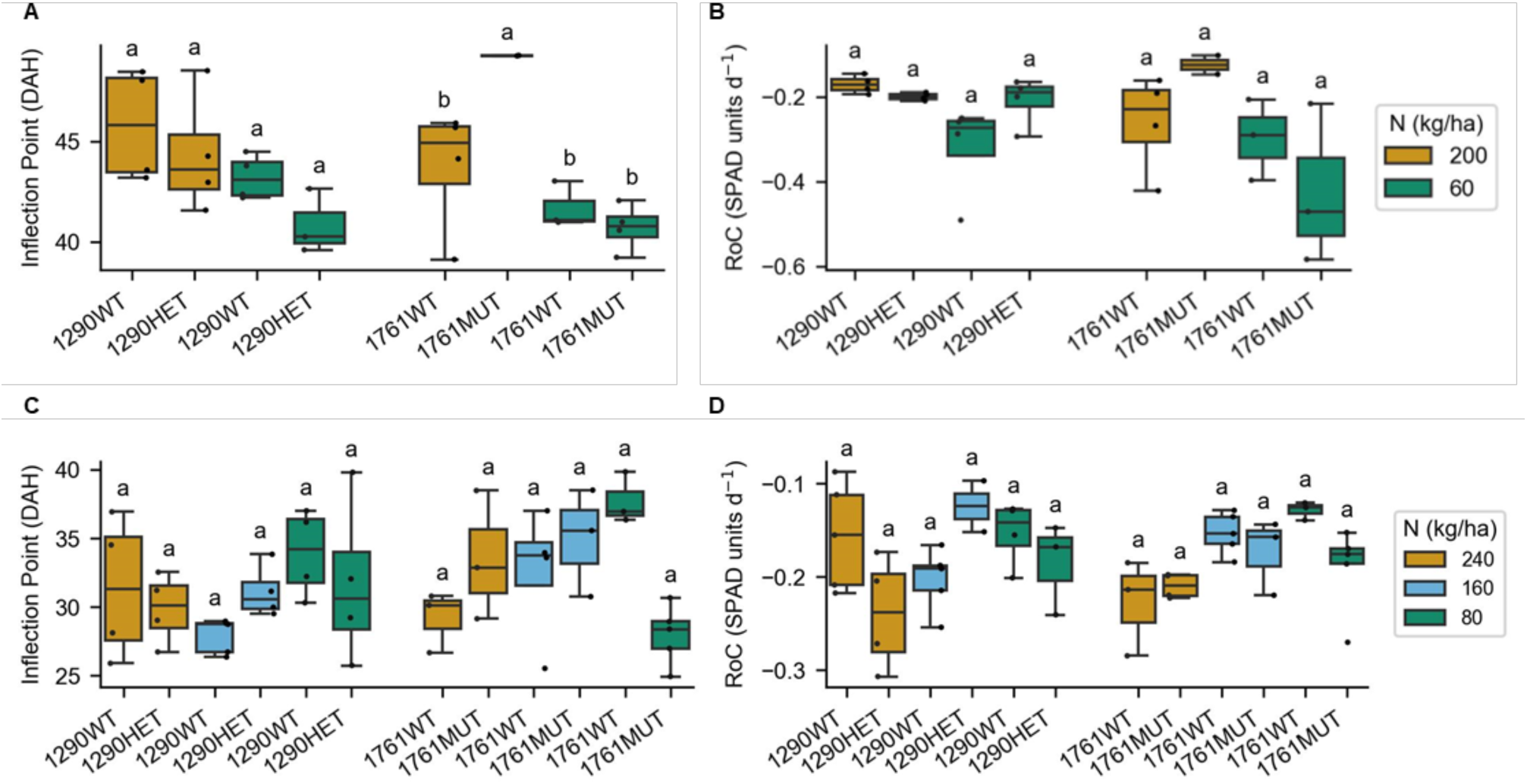
Effect of miR165/166-resistant *HB-2* alleles on flag leaf senescence. 1761MUT took more days to reach the inflection point compared to 1761WT under high N whereas no difference is observed in *CAD1290* **(A)** or for rate of change in either family **(B)** in University of Nottingham experiment. Similar results for inflection point **(C)** and rate of change **(D)** were observed in University of Adelaide experiment. Letters show significant differences based on Fisher’s LSD (α = 0.05) test with Benjamini-Hochberg adjusted p value to control the false discovery rate.

Because senescence is a key process associated with protein accumulation in wheat grain, these findings were further confirmed in Expt. 2. The validation experiment was conducted under controlled conditions (16 h light and 22 °C/15 °C (day/night)). The results were consistent with the previous experiment, with no difference in *CAD1290* genotypes, whereas 1761MT again showed delayed (but non-significant) senescence under both high and low N **(Fig. 1C-D; Supplementary Table 6)**. This analysis confirmed that the miR165/166 resistant *HB-2* alleles did not accelerate senescence, indicating GPC improvements associated with *HB-2* are unlikely to compromise photosynthetic duration or yield.

To test the effect of *HB-2* alleles on photosynthesis, we measured the operating efficiency of PSII (Fv’/Fm’), transpiration rate (mmol H_2_0 m^−2^ s^−1^) and leaf temperature (°C) in Expt. 1 through the Li-Cor 6400 photosynthesis system. Data were collected one and three weeks after heading and similar results were observed. Fv’/Fm’ was significantly higher in 1761MT compared to 1761WT under both N conditions meaning the mutant was able to use a greater portion of the absorbed light in PSII, potentially driving more photosynthesis **(Supplementary Fig. 8A)**. Transpiration rate was higher in both genotypes with miR165/166 resistant *HB-2* alleles, but the difference was not significant **(Supplementary Fig. 8B)**. This could indicate more CO_2_ uptake and better photosynthetic efficiency in the mutants but also increased water loss. However, there was no difference in leaf temperature, indicating that the difference in PSII efficiency was due to physiological difference in photosynthetic capacity or stomatal regulation, rather than due to thermal stress **(Supplementary Fig. 8C)**. Overall, the combined results from senescence and photosynthesis analyses suggest that senescence was not accelerated by miR165/166 resistant *HB-2* alleles and photosynthesis was positively affected, particularly by the variant *HB-A2* allele.

### *HB-2* alleles affect spike architecture under N limitation

To test if the increased expression of *HB-2* leads to other modifications beyond secondary spikelets, we measured spike and plant architecture traits in Expt. 1. Spike length was moderately reduced under low N in 1761WT but not in 1761MT. Similarly, rachis length was reduced moderately (non-significant) in 1290WT compared to 1290HT and significantly in 1761WT compared to 1761MT (G × N, *p* 0.0402*) under low N. Under optimum N conditions, both the spike and rachis of WT tended to be longer than those of mutants, although the difference was not significant. Similar observations were observed for spike area, which was reduced under low N in 1290WT, but not in 1290HT (G × N *p* 0.0102*) **(Supplementary Table 7)**. Overall, apart from the development of secondary SpS, spike length, rachis length and spike area were resistant to reductions in the mutants under N limitation. These results indicate that the miR165/166 resistant *HB-2* alleles may not influence the spike architecture under optimum N but induces tolerance of N limitation, making the plants more adaptable to environmental variation. Greater spike length, rachis length and spike area could be associated with improved spike vascular connections, which would not only improve the assimilation of nutrients under low N but also reduce heterogeneity along the spike **(Slafer *et al*., 2022)**.

### *HB-2* alleles enhance N transport and use efficiency

We hypothesised the architectural modifications by *HB-2* improve N uptake and translocation to grain in plants under N limitation. This was tested in Expt. 1 by investigating N accumulation at anthesis and harvest, N remobilisation and use efficiencies. At anthesis, a significant genotype effect (*p* 0.0159*) was observed for *CAD1761* with more AGN (mg) uptake per main shoot by 1761MT compared to 1761WT under both optimum N (63.13±8.77 mg vs. 54.34±3.52 mg) and low N (51.59±5.94 mg vs. 41.01±4.98 mg). No significant effect was observed on AGN at harvest, possibly due to high variability among replicates. Nonetheless, the average AGN reduction per main shoot in 1761WT compared to 1761MT was much larger – 19.85 mg N reduction in WT compared to 6.08 mg in MT under low N **(Table 2)**. This indicated that 1761MT was more resistant to N and yield losses under N limitation, which complemented the findings on spike architecture. Post-anthesis N uptake (PANU) was not affected by genetic or N differences, or G × N interaction **(Table 2)**. These results suggest that protein increase by *HB-2* alleles was associated with pre-anthesis N uptake and better mobilisation of already available AGN at anthesis in the vegetative organs. N partitioning index (NPI) of plant components at harvest showed interesting differences with higher NPI in stem (*p* 0.0134*) and chaff (*p* 0.0633) and less in grain (*p* 0.0461*) of 1290HT compared to the WT under both N levels at harvest **(Table 2)**. This suggests that although 1290HT has an apparently higher GPC at harvest (see **Table 1**), there may still be potential to transfer more N from vegetative tissue to the grain. No differences were observed in other plant-component NPIs as well as N-remobilisation efficiency for either 1761MT or 1290HT compared to the wild types. N-uptake efficiency and N-use efficiency significantly increased under low N across genotypes (*p* <0.001*** for both), whereas no effect was observed on N-utilisation efficiency. Moreover, significant G × N effect was observed in *CAD1761* for NUpE (*p* 0.0131*) and NUE (*p* 0.0456*) with the mutant showing relatively greater increase for both traits under low N compared to the wild type **(Table 2)**.

**Table 2:** Split-plot analysis of variance for N partitioning indices and N-use ejiciency in experiment 1. Values in table represent mean ± standard deviation (n = 4)

| Genotype | High N | Low N | Difference | ANOVA |
| --- | --- | --- | --- | --- |
| **Above ground N (mg) at anthesis** |  |  |  |  |
| 1290W | 61.08±8.32 | 46.01±6.28 | 15.07 | N: $p = 0.003^{**}$ |
| 1290H | 55.12±5.09 | 40.44±3.82 | 14.67 | G: $p = 0.2134$ |
| | | | | $G \times N$: $p = 0.9507$ |
| 1761W | 54.34±3.52 | 41.01±4.98 | 13.33 | N: $p = 0.0046^{**}$ |
| 1761M | 63.13±8.77 | 51.59±5.94 | 11.54 | G: $p = 0.0159^*$ |
| | | | | $G \times N$: $p = 0.763$ |
| **Leaf N partitioning index (%) at anthesis** |  |  |  |  |
| 1290W | 40.37±2.17 | 39.34±5.1 | 1.03 | N: $p = 0.2925$ |
| 1290H | 38.52±7.06 | 32.86±2.25 | 5.66 | G: $p = 0.142$ |
| | | | | $G \times N$: $p = 0.4545$ |
| 1761W | 38.99±3.43 | 34.62±2.49 | 4.37 | N: $p = 0.0466^*$ |
| 1761M | 38.6±4.6 | 33.61±2.54 | 4.99 | G: $p = 0.7128$ |
| | | | | $G \times N$: $p = 0.8734$ |
| **Stem N partitioning index (%) at anthesis** |  |  |  |  |
| 1290W | 31.04±3.89 | 32.45±7.64 | -1.41 | N: $p = 0.8275$ |
| 1290H | 36.93±6.39 | 36.76±2.3 | 0.17 | G: $p = 0.2354$ |
| | | | | $G \times N$: $p = 0.7811$ |
| 1761W | 31.45±2.94 | 33.27±2.86 | -1.83 | N: $p = 0.5303$ |
| 1761M | 31.26±3.94 | 32.08±3.86 | -0.82 | G: $p = 0.5333$ |
| | | | | $G \times N$: $p = 0.8096$ |
| **Spike N partitioning index (%) at anthesis** |  |  |  |  |
| 1290W | 28.6±4.88 | 28.21±4.51 | 0.39 | N: $p = 0.1846$ |
| 1290H | 24.55±3.07 | 30.38±1.46 | -5.83 | G: $p = 0.5846$ |
| | | | | $G \times N$: $p = 0.1377$ |
| 1761W | 29.57±3.72 | 32.11±4.39 | -2.54 | N: $p = 0.3337$ |
| 1761M | 30.15±5.22 | 34.31±6.18 | -4.16 | G: $p = 0.4452$ |
| | | | | $G \times N$: $p = 0.8072$ |
| **Above ground N (mg) at harvest** |  |  |  |  |
| 1290W | 73.39±11.08 | 63.36±11.25 | 10.03 | N: p = 0.1153 |
| 1290H | 62.27±12.38 | 52.12±7.56 | 10.14 | G: p = 0.1516 |
|  |  |  |  | G×N: p = 0.9922 |
| 1761W | 69.5±20.64 | 49.65±6.25 | 19.85 | N: p = 0.05. |
| 1761M | 67.72±16.42 | 61.64±7.14 | 6.08 | G: p = 0.6916 |
|  |  |  |  | G×N: p = 0.2416 |
| **Post anthesis N uptake (mg)** |  |  |  |  |
| 1290W | 12.31±15.79 | 17.35±14.78 | -5.04 | N: p = 0.5662 |
| 1290H | 7.15±14.54 | 11.68±10.3 | -4.53 | G: p = 0.3806 |
|  |  |  |  | G×N: p = 0.9752 |
| 1761W | 15.16±23.72 | 8.64±9.19 | 6.52 | N: p = 0.9241 |
| 1761M | 4.59±14.3 | 10.05±9.81 | -5.46 | G: p = 0.7269 |
|  |  |  |  | G×N: p = 0.3045 |
| **Leaf N partitioning index (%) at harvest** |  |  |  |  |
| 1290W | 5.85±1.93 | 5.26±0.83 | 0.59 | N: p = 0.5543 |
| 1290H | 5.48±1.71 | 5.23±1.24 | 0.24 | G: p = 0.8527 |
|  |  |  |  | G×N: p = 0.8014 |
| 1761W | 6.76±1.24 | 4.24±1.68 | 2.52 | N: p = 0.0163* |
| 1761M | 7.92±3.72 | 5.59±1.25 | 2.33 | G: p = 0.5421 |
|  |  |  |  | G×N: p = 0.9048 |
| **Stem N partitioning index (%) at harvest** |  |  |  |  |
| 1290W | 6.77±0.84 | 7.07±1.42 | -0.29 | N: p = 0.6563 |
| 1290H | 8.87±1.27 | 9.28±2.2 | -0.42 | G: p = 0.0134* |
|  |  |  |  | G×N: p = 0.9363 |
| 1761W | 7.53±1.92 | 8.25±1.22 | -0.73 | N: p = 0.7085 |
| 1761M | 10.15±5.02 | 8.15±1.78 | 2 | G: p = 0.4811 |
|  |  |  |  | G×N: p = 0.433 |
| **Chaff N partitioning index (%) at harvest** |  |  |  |  |
| 1290W | 3.62±1.02 | 2.93±0.34 | 0.7 | N: p = 0.1059 |
| 1290H | 4.47±0.36 | 4.03±0.43 | 0.44 | G: p = 0.0633. |
|  |  |  |  | G×N: p = 0.6827 |
| 1761W | 3.34±0.48 | 3.4±0.57 | -0.05 | N: p = 0.363 |
| 1761M | 4.09±1.28 | 3.3±0.36 | 0.79 | G: p = 0.5688 |
|  |  |  |  | G×N: p = 0.3014 |
| **Rachis N partitioning index (%) at harvest** |  |  |  |  |
| 1290W | 1.17±0.26 | 0.83±0.14 | 0.35 | N: p = 0.0269* |
| 1290H | 1.17±0.09 | 0.93±0.05 | 0.24 | G: p = 0.1727 |
|  |  |  |  | G×N: p = 0.5942 |
| 1761W | 1.13±0.47 | 0.98±0.19 | 0.15 | N: p = 0.1446 |
| 1761M | 1.22±0.45 | 0.83±0.15 | 0.39 | G: p = 0.9231 |
|  |  |  |  | G×N: p = 0.4784 |
| **Grain N partitioning index (%) at harvest** |  |  |  |  |
| 1290W | 82.57±2.33 | 83.92±1.92 | -1.34 | N: p = 0.4313 |
| 1290H | 80.02±1.5 | 80.51±1.5 | -0.5 | G: p = 0.0461* |
|  |  |  |  | G×N: p = 0.7121 |
| 1761W | 81.25±3.47 | 83.13±2.7 | -1.88 | N: p = 0.2008 |
| 1761M | 76.62±10.07 | 82.13±2.96 | -5.51 | G: p = 0.5408 |
|  |  |  |  | G×N: p = 0.5069 |
| **Leaf N remobilisation (%)** |  |  |  |  |
| 1290W | 82.98±4.52 | 81.36±4.36 | 1.62 | N: p = 0.0871. |
| 1290H | 84.28±3.38 | 79.03±7.37 | 5.26 | G: p = 0.8258<br>G×N: p = 0.3223 |
| 1761W | 77.82±6.47 | 85.1±7.04 | -7.28 | N: p = 0.1728 |
| 1761M | 79.3±4.59 | 80.4±1.91 | -1.1 | G: p = 0.5178<br>G×N: p = 0.2972 |
| **Stem N remobilisation (%)** |  |  |  |  |
| 1290W | 73.44±5.16 | 67.87±14.33 | 5.57 | N: p = 0.3298 |
| 1290H | 72.16±8.28 | 67.49±8.02 | 4.67 | G: p = 0.8671<br>G×N: p = 0.9287 |
| 1761W | 70.38±6.58 | 69.13±8.26 | 1.25 | N: p = 0.8836 |
| 1761M | 66.61±13.69 | 69.52±5.81 | -2.91 | G: p = 0.5742<br>G×N: p = 0.7146 |
| **N remobilisation efficiency (%)** |  |  |  |  |
| 1290W | 79.05±3.22 | 77.74±4.69 | 1.31 | N: p = 0.3762 |
| 1290H | 77.36±4.65 | 74.74±4.49 | 2.62 | G: p = 0.2506<br>G×N: p = 0.7611 |
| 1761W | 76.55±5.03 | 79.21±5.84 | -2.66 | N: p = 0.371 |
| 1761M | 76.42±5.4 | 78.73±2.9 | -2.31 | G: p = 0.8624<br>G×N: p = 0.9474 |
| **N uptake efficiency (%)** |  |  |  |  |
| 1290W | 20.76±3.14 | 59.75±10.61 | -38.99 | N: p = 1e-04*** |
| 1290H | 17.62±3.5 | 49.16±7.13 | -31.54 | G: p = 0.1062<br>G×N: p = 0.3226 |
| 1761W | 19.66±5.84 | 46.83±5.89 | -27.16 | N: p = <0.001*** |
| 1761M | 19.16±4.65 | 58.13±6.73 | -38.97 | G: p = 0.3745<br>G×N: p = 0.0131* |
| **N utilisation efficiency (g GY / g N)** |  |  |  |  |
| 1290W | 50.51±1.57 | 51.26±2.87 | -0.75 | N: p = 0.5094 |
| 1290H | 51.96±5.86 | 53.82±2.44 | -1.85 | G: p = 0.215<br>G×N: p = 0.7752 |
| 1761W | 45.16±8.38 | 49.7±4.89 | -4.54 | N: p = 0.1993 |
| 1761M | 45.15±6.48 | 48.54±1.68 | -3.38 | G: p = 0.7857<br>G×N: p = 0.8402 |
| **N use efficiency (g GY / g N)** |  |  |  |  |
| 1290W | 10.51±1.77 | 30.51±4.85 | -20 | N: p = 1e-04*** |
| 1290H | 9.29±2.77 | 26.55±4.83 | -17.25 | G: p = 0.173<br>G×N: p = 0.4965 |
| 1761W | 8.54±0.81 | 23.2±3.15 | -14.66 | N: p = <0.001*** |
| 1761M | 8.85±2.97 | 28.25±3.71 | -19.4 | G: p = 0.3597<br>G×N: p = 0.0456* |

Significance codes: 0 '\*\*\*' 0.001 '\*\*' 0.01 '\*' 0.05

Overall, these results suggest the miR165/166 resistant alleles of *HB-2* improve a plant’s potential to uptake total N pre-anthesis which may contribute to protein production during grain maturation. The effect of *HB-2* alleles on genetic factors during grain maturation remains to be seen, which could be investigated through transcriptomic analyses of the developing grain. Comparatively, 1761MT looks more promising than 1290HT with higher tolerance of N limitation in terms of better AGN accumulation at harvest and balanced NPI as well as improved NUE.

### *HB-2* expression is enhanced in developing spike under N limitation

Dixon *et al*. (2022) reported that the modifications in vascular development and GPC increase associated with the miR165/166 resistant *HB-2* alleles replicated phenotypes observed in wild-type wheat grown under high N. Based on their proposal, we sought to investigate the contribution of *HB-2* to GPC in wild-type plants by analysing its response to N. We hypothesised that *HB-2* expression would increase in a wild-type plant under high N and lead to increased grain protein. An alternate hypothesis, which is supported by our glasshouse experiments, is that *HB-2* expression increases under N limitation, inducing changes in spike architecture and influencing N remobilisation. As spikelet architecture is determined during early developmental stages, *HB-2* expression was investigated in developing spikes, stem, and leaf tissue at lemma primordium and terminal spikelet stages under three N levels. Relative expression levels of all three *HB-2* homeologues (*HB-A2*, *HB-B2* and *HB-D2*) were analysed using a mixed effect model with individual and all interaction terms followed by a less complex model with a maximum of two interaction terms. Both models indicated a strong effect of tissue for all homeologues, and there was no interaction effect with N level across tissues or stages. The convergence of both models failed, which indicates multi-level complexity compared to the amount of data or multicollinearity among predictor variables. Hence, a one-way ANOVA was considered to analyse the N effect at each stage and tissue. This showed a significant response to N levels by *HB-A2* (*p* 0.0024**) and *HB-D2* (*p* 0.014*) in the spike at the lemma primordium stage and *HB-D2* (*p* 0.0456*) in the spike at the terminal spikelet stage. *HB-B2* showed a significant response to N level in stem at terminal spikelet stage (*p* 0.0223*); no response was observed in leaf tissues **(Supplementary Table 8)**. After the gene’s response to N levels in spike and stem was confirmed, post-hoc analysis was conducted at different stages using Tukey’s HSD method. At lemma primordium, all homeologues showed a significant increase in the spike under low N **(Fig. 2A)**, whereas no difference was observed in the stem **(Fig. 2B)**. It must be noted that high variability among replicates was observed; nonetheless, at least two homeologues showed a similar trend. The increased *HB-2* expression in the spike may be responsible for the increased spike and rachis length under reduced N (see **Supplementary Table 7**). Together, these results support the second hypothesis and suggest that plants respond to early N limitation by increasing the expression of *HB-2* in the developing spike, which in turn induces changes in the spike architecture. The mutant lines that have inherently higher expression of *HB-2* develop longer spikes under reduced N that results in better N accumulation and distribution at later stages. At terminal spikelet, *HB-D2* was the only homeologue to display significantly higher expression in spike at low N, whereas the other copies did not show significant differences **(Fig. 2C)**. In stem tissue at terminal spikelet, the expression of *HB-B2* increased significantly under the high N treatment. *HB-A2* and *HB-D2* expression showed a similar trend but there was high variability within replicates **(Fig. 2D)**. These results suggest *HB-2* shows a stage and tissue specific response to N application.

**Figure 2:**
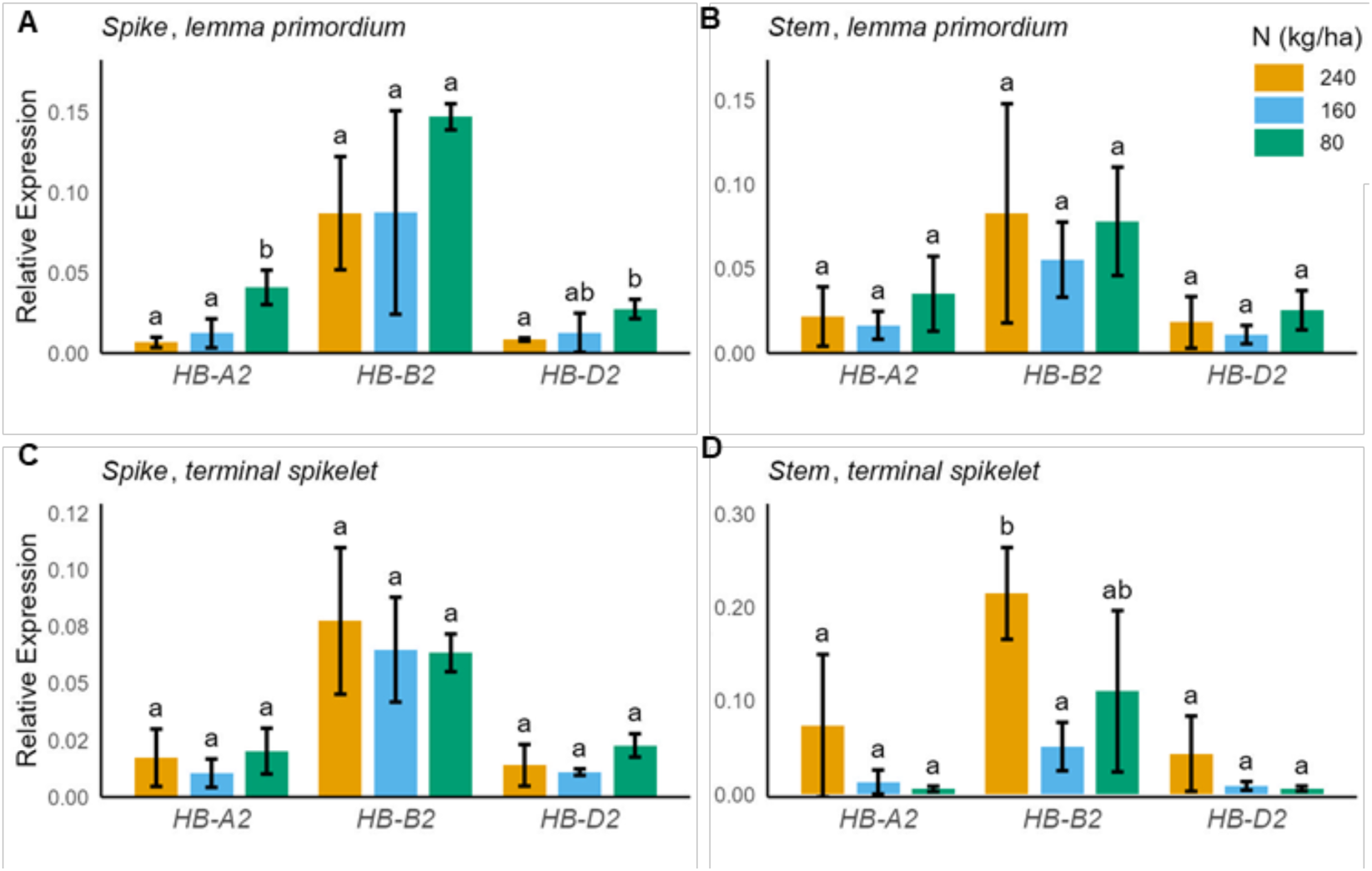
*HB-2* expression under different N levels during early development. *HB-A2* and *HB-D2* expressions were significantly high under low N in the developing spike **(A)** whereas no response to N levels was observed in stem **(B)** at lemma primordium stage. At terminal spikelet, no differences in expression were observed in spike under different N levels **(C)** whereas all homeologues showed an increasing expression trend in stem under high N, with *HB-B2* being significantly high **(D)**. Asterisks show significance level (’***’ 0.001 ‘**’ 0.01 ‘*’ 0.05) from one-way ANOVA. Bars show mean ± SD (n = 3). Letters show significant differences based on Tukey’s HSD (α = 0.05) with adjusted p value to control the family-wise error rate.

Based on the increased *HB-2* expression in developing spike at lemma primordium and terminal spikelet, we predicted that the higher GPC is facilitated by the spike architectural changes that occur during early spike development, rather than higher *HB-2* expression in the grain directly influencing protein levels. To test this further, we analysed the expression of *HB-2* homeologues in developing grain at 15, 23 and 30 days after anthesis using RNA-Seq. All *HB-2* homeologues were lowly expressed in the developing grain under both high N and low N **(Supplementary Fig. 9)**, supporting our hypothesis. These findings open a question for future research: do miR165/166-resistant *HB-2* alleles improve GPC by enhancing source remobilisation or by generating a higher sink strength, or a combination of both?

### Introgression of *HB-2* alleles into cultivated elite wheat improves GPC

To test if the miR165/166 resistant *HB-2* alleles improve GPC in the background of elite wheat cultivars grown by farmers, we introgressed miRNA-resistant *HB-A2* and *HB-D2* alleles into three Australian elite cultivars, Mace, Rockstar and Sherrif. These genotypes were compared to segregating wild-type sibling lines. GPC and yield component traits were analysed at two field experimental sites: the first near the Roseworthy Campus of University of Adelaide (Trial 1) and the second at a nearby site managed by Australian Grain Technologies (Trial 2). At Trial 1, GPC was significantly higher in Mace/1761MT and Rockstar/1761MT compared to their wild-type siblings, but no effect was observed in Sherrif/1761MT **(Fig. 3A)**. Regarding yield-component traits, TGW was significantly lower in Mace/1761MT but GN per spike was higher, which meant the GW per spike was not affected; no effect in yield components was observed in the Rockstar background whereas both TGW and GW per spike increased in Sherrif/1761MT compared to Sherrif/1761WT **(Fig. 3B-D)**. These results showed that an increase in GPC by introgressing *HB-A2* beneficial allele was achieved without a yield penalty. At Trial 2, GPC was significantly higher in Mace/1761MT and moderately (non-significant) higher in Rockstar/1761MT compared to their wild-type siblings **(Fig. 3E)**. All the studied yield components (except TGW in Sherrif/1761MT) were significantly reduced in both backgrounds compared to their wild-type siblings, indicating that environment and soil conditions may have impacted the performance of the *HB-A2* alleles **(Fig. 3F-H)**. To check if there was a G × E interaction, we performed a two-way ANOVA for GPC by including the interaction term in the model, which yielded a non-significant *p* value **(Supplementary Table 9)**. This indicates that the positive effect of *HB-A2* on GPC is independent of environment, but its effect on yield components may vary in different growth conditions.

**Figure 3:**
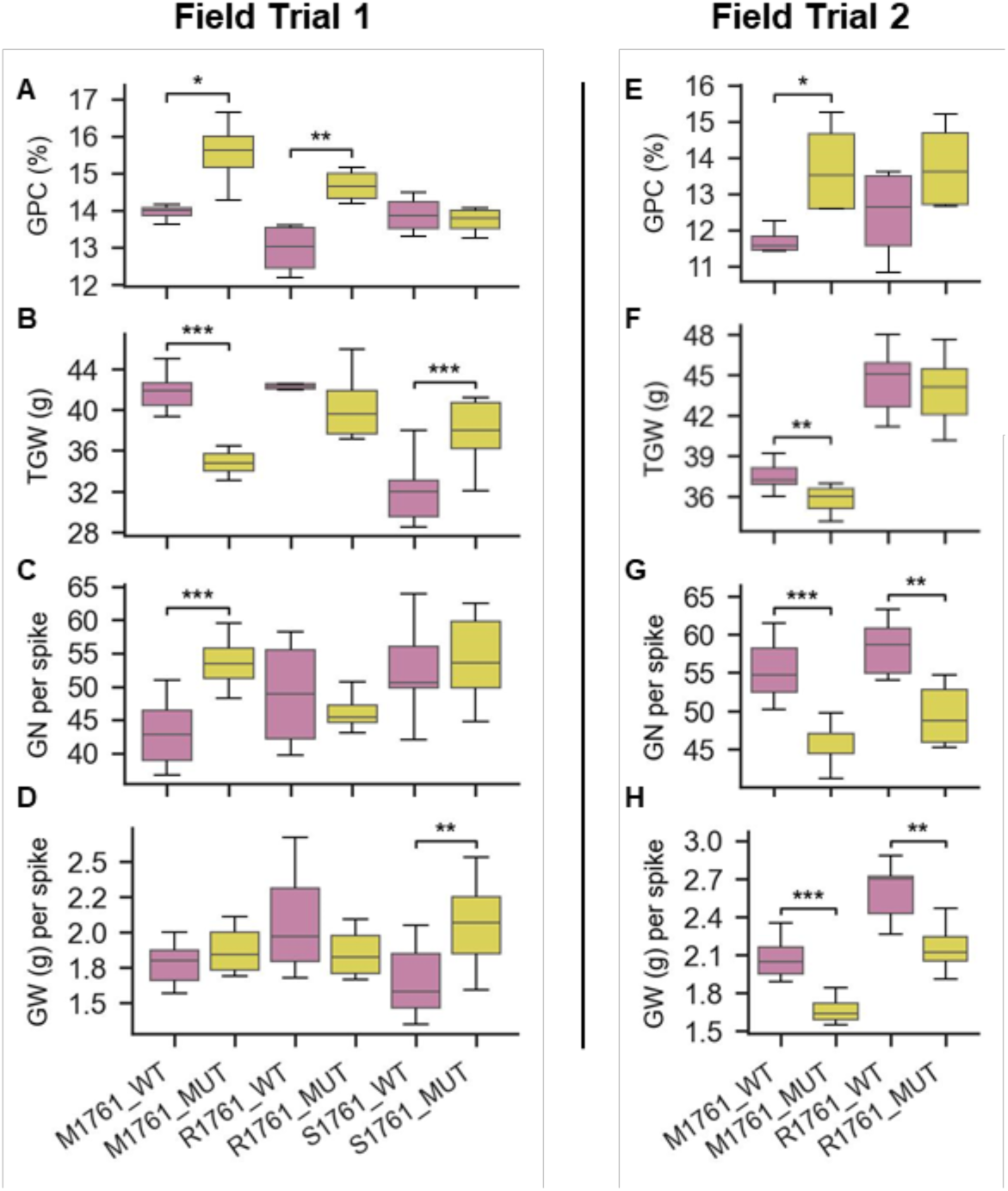
Effect of miR165/166-resistant *HB-A2* on GPC and yield components in elite wheat genetic background. GPC, GN per spike, GW (g) per spike and TGW (g) were analysed in elite wheat background by crossing 1761WT and 1761MT with Mace (M-1761), Rockstar (R-1761) and Sherrif (S-1761) at two field trials, University of Adelaide Roseworthy Campus **(A-D)** and Australian Grain Technologies **(E-H)**. All analyses were conducted by independent samples *t* test to test significant differences in means (n = 4 for GPC and n =10 for yield traits). Asterisks represent *p* values at * <0.05, ** <0.01 and *** <0.001.

In the case of *HB-D2*, GPC and yield parameters were analysed in all genotypes at the two sites except for those in the background of Sheriff, where only GPC was measured because these lines flowered late. At Trial 1, GPC was significantly higher in Mace/1290MT, Rockstar/1290MT, Sherrif/1290HT and Sherrif/1290MT and a moderate (non-significant) increase was observed in Mace/1290HT and Rockstar/1290HT compared to their wild-type siblings **(Fig. 4A)**. GN and GW per spike were not affected in Mace and Rockstar with 1290HT alleles but reduced in genotypes carrying 1290MT alleles; TGW was reduced in Mace/1290HT **(Fig. 4B-D)**. Similar results were noted at Trial 2, that is significant increase GPC in Mace/1290MT and Rockstar/1290MT, and moderate (non-significant) increase in Mace/1290HT and Mace/1290MT compared to the wild types **(Fig. 4E)**. Furthermore, the yield components were not reduced in the heterozygous or mutant genotypes with Mace background; in fact, GN and GW (g) per spike were significantly higher in Mace/1290HT compared to Mace/1290WT, whereas all three yield traits showed a significant decrease in heterozygous and mutant genotypes with Rockstar background **(Fig. 4F-H)**. Once again, no G × E interaction was observed for GPC **(Supplementary Table 9)**, but there may be an environment specific yield performance. The higher GPC in elite wheat background carrying the miRNA-resistant *HB-2* alleles supports their introgression into breeding programs that aim to improve grain quality through enhanced protein content without significantly hampering yield. Future research would benefit from a broader analysis of G × E interaction on yield components including yield t/ha, senescence, NUE and other agronomically important traits.

**Figure 4:**
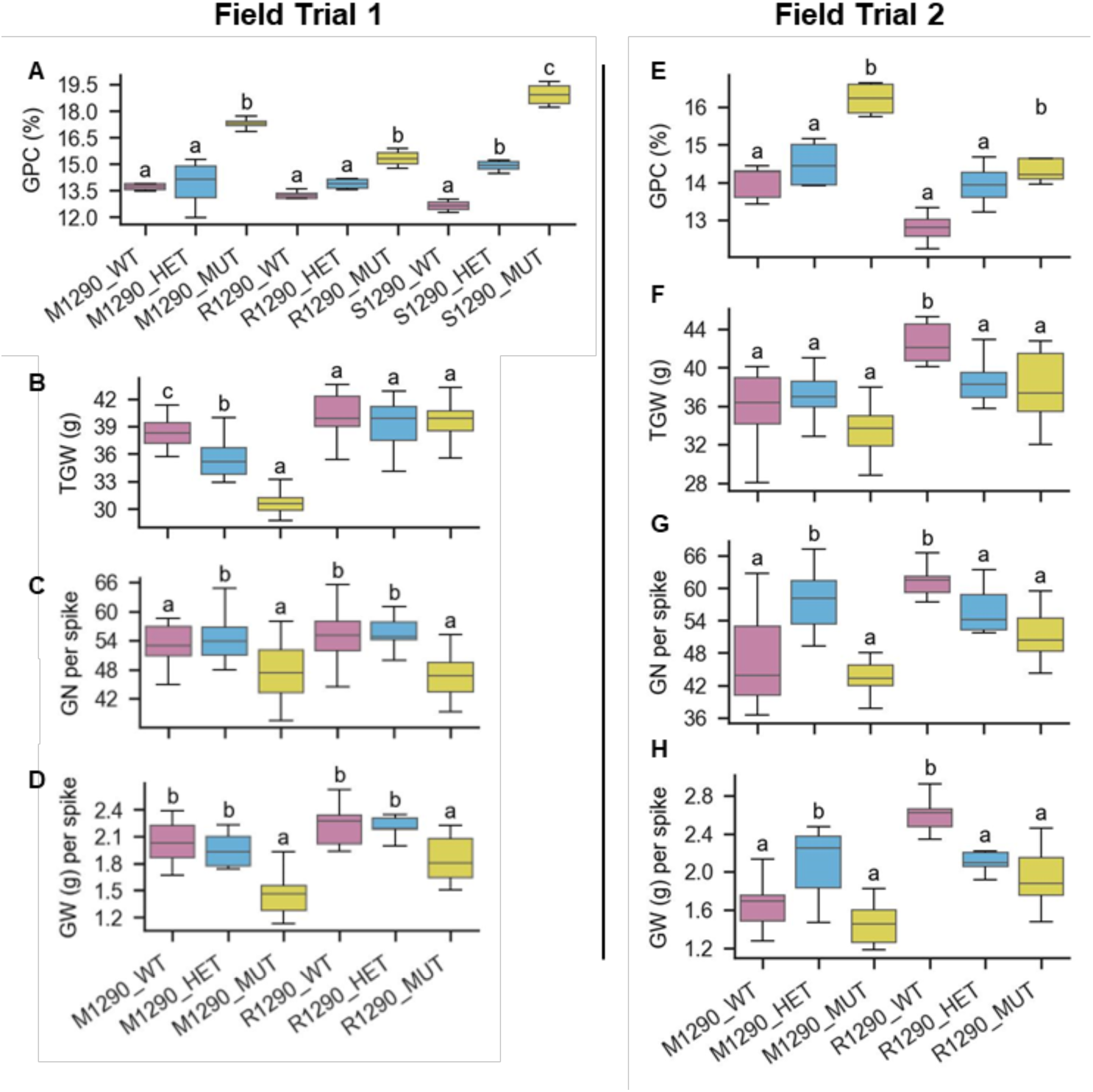
Effect of miR165/166-resistant *HB-D2* on GPC and yield components in elite wheat genetic background. GPC, GN per spike, GW (g) per spike and TGW (g) were analysed in elite wheat background by crossing 1290WT, 1290HT and 1290MT with Mace (M-1290), Rockstar (R-1290) and Sherrif (S-1290) at two field trials, University of Adelaide Roseworthy Campus **(A-D)** and Australian Grain Technologies **(E-H)**. All analyses were conducted by one-way ANOVA. Letters indicate significant differences in means (n = 4 for GPC and n = 10 for yield traits) as analysed by Tukey’s HSD.

## DISCUSSION

Wheat grains with more protein are commercially valuable because protein levels strongly influence bread quality. Recently, miR165/166 resistant alleles of *HB-2* were reported to improve protein content in wheat grains by introducing changes in plant vasculature, leading to enhanced remobilisation of N-based assimilates to the spike **(Dixon *et al*., 2022)**. Here, we asked if the miRNA-resistant *HB-2* alleles affect senescence and N uptake and/or utilisation components, and if they showed a yield trade-off as is often found in the case of GPC-enhancing loci such as *Gpc-B1* **(Tabbita *et al*., 2017)**. Our analysis of genotypes with wild-type and miRNA-resistant *HB-2* alleles under different N levels revealed that the increase in protein was not dependent on N supply, neither was it achieved at the expense of grain yield. The introgression of *HB-2* alleles into elite wheat cultivars confirmed that protein increase was not associated with reduced yield. In some cases, GW was reduced but the complimentary increase in GN per spike meant that yield remained stable. These results support introducing *HB-2* beneficial alleles into cultivars for GPC improvement.

There is a longstanding debate on whether genetic variation in grain protein is achieved through N source related to accumulation and/or remobilisation or altered N sink strength of the grains. Previous research reports improvement of GPC by remobilising more N to the grain, either through accelerated senescence by *Gpc-B1* functional alleles **(Uauy *et al*., 2006b)**, or by promoting development of more vascular bundles in the stems **(Dixon *et al*., 2022)**. Our results showed that expression of *HB-2* homeologues increased in spikes in response to N limitation during early inflorescence development. In combination with the previous analyses of *HB-2*, this result suggests that the higher GPC is a consequence of improved sink strength effected during early spike development. This model is supported by the limited expression of *HB-2* homeologues during grain development, and the reported role of HD-ZIP family transcription factors in inflorescence development, including the development of supernumerary spikelets **(Ariel *et al*., 2007; Dixon *et al*., 2022)**. Future research should further investigate the N source vs. sink relationship, and the role of HD-ZIP III family genes in regulating protein content in cereals and its relationship with yield. The role of HD-ZIP genes in improving grain yield through increase GN per spike has been reported for other genes as well, e.g., *GNI-A1* **(Sakuma *et al*., 2024)**. This presents a great opportunity to improve GPC in wheat without the expected yield trade-off.

Our results demonstrated that the improved GPC by higher *HB-2* expression was not caused by accelerated senescence, unlike *Gpc-B1* (see **Figure 1**). Nutrient remobilisation from vegetative to reproductive organs lies at the core of senescence, which also determines nutrient-use efficiency in cereals. This places senescence at the epicentre of grain yield and grain nutrient balance – referred to as “the dilemma of senescence” in a previous review **(Gregersen, 2011)**. On the one hand, accelerated senescence can improve N-harvest index and enhance protein content by reallocating more N to the grain **(Hirel *et al*., 2007; Uauy *et al*., 2006a; Uauy *et al*., 2006b; Waters *et al*., 2009)**; while on the other hand, delayed senescence can promote stay-green phenotypes insofar that it increases the grain filling period and photosynthesis duration, consequently improving yield **(Spano *et al*., 2003; Yang and Zhang, 2006)**. Therefore, identifying alternate processes that improve GPC without accelerating senescence or shortening grain filling would be desirable for breeding. Taken together with recent reports on the role of HD-ZIP proteins affecting grain yield and quality through influencing plant vascular and inflorescence architecture, our results suggest a complimentary approach towards protein improvement **(Dixon *et al*., 2022; Sakuma *et al*., 2024)**. Therefore, *HB-2* would be a promising candidate for introgression into cultivated bread-making wheat for investigating the protein-yield dilemma in different environments. Investigating the interaction between genes regulating vascular and inflorescence architecture and whole-plant senescence would further benefit grain quality research.

## CONCLUSION

These results support the hypothesis that miR165/166-resistant *HB-2* alleles improve GPC without significantly compromising grain yield. Mutations in the miR165/166 binding site of *HB-2* do not accelerate leaf senescence or negatively impact photosynthetic efficiency of the plants; hence, yield is unlikely to be affected, as demonstrated in the *HB-2* introgression lines. These results also emphasise the complexity of GPC and its interaction with multiple genetic and environmental factors. Hence, efforts to identify more promising candidate GPC alleles— resistant to environmental fluctuations—must continue **(Safdar *et al*., 2023b)**. Advancements in next generation sequencing and machine learning could be combined to investigate genetically rich germplasm for the discovery of new beneficial alleles.

## ACKNOWLEDGEMENTS

This work was supported by the Australian Research Council (FT210100810), the Biotechnology and Biological Sciences Research Council (BBSRC; BB/V018108/1, BB/W006979/1, BB/V004115/1), and the Adelaide–Nottingham Alliance for Joint PhD. RAB acknowledges BBSRC funding through a Discovery Research Fellowship (BB/S011102/1), New Investigator Grant (BB/X014843/1), and DSW Partner Grant (BB/X018806/1). We thank Mark Meacham, Catherine (University of Nottingham), and Lidia Mischis (University of Adelaide) for assistance with glasshouse experiments; Elizabeth Goodman (University of Nottingham) for phenotype data collection; Yue Qu (University of Adelaide) for KASP analysis; Alix Cornish (Campden BRI) for Kjeldahl N analysis; and Alistair Pearce (University of Adelaide), Cathrine Ingvordsen, James Walter and Takara Fort (Australian Grain Technologies) for field trial support and GPC analyses.

## DATA AVAILABILITY

All phenotypic data used in these experiments are available in supplemental materials. RNA-Seq data used to test the expression levels of *HB-2* homeologues in developing grains come from an ongoing project at Boden Lab and will be released soon. Questions related to RNA-Seq data can be addressed to Dr Scott Boden.

## AUTHOR CONTRIBUTIONS

SAB, MJF, IDF, RAB and IRS conceived and supervised the project. LBS, SAB, MJF, MP and AL conducted the experiments. LBS analysed the data with inputs from SAB and MJF and wrote the initial draft of manuscript. IDF, SAB, MJF and RAB reviewed and edited the manuscript. All authors read and approved the final version.

## COMPETING INTERESTS

The authors declare no competing interests.

## Notes

### Competing Interest Statement

The authors have declared no competing interest.

